# Comprehensive Characterization of the Promoter Proximal Proteome of Single Copy Locus *FOXP2*

**DOI:** 10.1101/2025.07.10.663086

**Authors:** Tim MG MacKenzie, Lucia Ramirez, Ruiqi Jian, Lihua Jiang, Michael P Snyder

## Abstract

Determining the proteins interacting with sequence-defined chromatin segments is a critical step in understanding gene expression and developing experimental interventions in the process. We used genetically targeted proximity labeling with dCas9-APEX2 to specifically biotinylate the promoter proximal proteome of the single copy locus *FOXP2* in live HEK293 cells. To identify labeled proteins in a discovery-based manner, we utilized quantitative mass spectrometry. Specifically, online 2D-LC coupled directly to a tribrid mass spectrometer to enable real-time database searching synchronous precursor selection MS3 provided deep proteome coverage and accurate quantitation via isobaric tandem mass tags. We inferred 6,039 proteins from our sample using Proteome Discoverer and performed bioinformatic analysis on quantified proteins to identify 373 significantly enriched proteins at the active promoter (Storey-*q*<.05, FC>1.2). These proteins were enriched for transcription factors and components of the spliceosome. To validate our candidate transcriptional regulators, we utilized computationally predicted transcription factor binding and the >200 ChIP-Seq experiments performed in HEK293 cells by ENCODE. In addition to validating several candidate transcription factors as binders of the targeted genomic locus, we identify IRF2BP2 as a negative regulator of *FOXP2* transcription.

## Introduction

The factors that determine gene expression are the central link between genotype and phenotype. In the canonical model of transcriptional regulation, genes contain proximal promoters and distal enhancers that contain specific DNA sequences recognized by transcription factors [1–3]. These transcription factors bind to DNA and recruit transcriptional machinery and chromatin remodeling proteins to impact gene expression. The dynamic composition of proteins at a promoter integrates environmental and cell-intrinsic signals to determine the activation or silencing of a gene at steady-state and in response to external cues. Determining the composition of proteins proximal to a gene promoter is a key step in building a mechanistic understanding of gene expression and developing experimental interventions in the process.

There is a growing appreciation for the role of alternative promoter selection in adding an additional layer of gene regulation [4–9]. Results from both long read [10] and traditional short read [11] sequencing have indicated that alternative promoters resulting in different transcription start sites (TSSs) and 5’ UTRs are the major driver of transcriptome diversity in many cancers and can provide a better patient prognosis than gene-level expression. The choice of TSS and the resulting 5’ UTR can have a meaningful impact on translation efficiency of the resulting gene, even when the same protein isoform is encoded [12–13]. Some promoters are ubiquitously active, while others are tissue, developmental time point, or disease-state specific [14]. Tumors in prostate cancer have been shown to increase alternative promoter usage as the disease progresses [15]. Efforts to identify promoter-proximal proteomes to understand the molecular mechanisms governing alternative promoter selection require tools that can detect minute analytes across a wide dynamic range and specific targeting to resolve a ∼1kbp promoter.

The isolation of sequence defined chromatin regions represents a major biochemical challenge due to two competing experimental realities: low intrinsic signal coupled to high background [16–17]. There are only two counts of a single copy locus per diploid cell, while a promoter on the kbp scale represents only ∼0.0001% of the Gbp sized human genome. Compounding the challenge is the biophysical similarity of chromatin - negatively charged DNA and positively charged nucleosomes. Furthermore, there is a huge dynamic range of proteins of interest. The copy number of transcription factors per animal cell is on the order of 10,000-300,000 per nucleus [18], while the winding of ∼150 bp of DNA around each nucleosome means there are tens of millions of histones per cell [19, 20]. If a promoter is a needle in a haystack, identifying the promoter-proximal proteome is akin to identifying the specific pieces of straw that surround the needle in the pile [21]. Early efforts to isolate sequence-defined chromatin regions focused on yeast to reduce background with its smaller genome [22–23] or telomeres to increase signal due to their abundance [24–25]. Recent efforts have utilized nuclease-dead dCas9-based strategies to target and isolate single copy loci in model organisms from human cells [26–27] to plants [28], relying on formaldehyde-crosslinking proteins to chromatin and isolating them. Complementary approaches have fused proximity labeling enzymes [29–30] to dCas9 to specifically biotinylate the proteins physically near the targeted promoter region [31–41]. These proximity labeling approaches preclude the need to cross-link chromatin, thereby enabling capture of chromatin associated proteins that are not covalently linked to DNA by formaldehyde, and allow for stringent wash steps using streptavidin-based purification of the labeled local proteome [42].

Mass spectrometry proteomics is the analytical tool used for proteomic profiling of chromatin [43]. While some approaches have compared the whole chromatin proteome between cell states [44–45], advances in protocols and instrumentation have enabled deep proteome coverage even on material-limited samples like single cells [46–47] and promoter-proximal proteomes. Highly accurate quantitation and multiplexing can be achieved with tandem mass tags (TMTs) that barcode individual samples before combining them together for LC-MS analysis [48]. A typical bottom-up data dependent acquisition shotgun proteomics approach digests the protein fraction of biological samples to the corresponding peptides which are separated by retention time, measured on the mass spectrometer, and fragmented to produce characteristic patterns that are also measured on a mass analyzer (MS2) [49–50]. Extensive fractionation using an online 2D-LC system allows for deeper proteome coverage by resolving complex chromatography peaks [51]. A third mass spectrometry step (MS3) to measure TMT signals ensures greater fidelity to the underlying ground truth quantitation by accounting for reporter ion ratio compression, while real time database searching and synchronous precursor selection (RTS-SPS) economizes on instrument time and increases signal, respectively [52–54]. The Orbitrap Ascend tribrid mass spectrometer design allows for deep proteome coverage using a TMT-based RTS-SPS-MS3 proteomics experiment [55–56].

The transcription factor *FOXP2* is among the most highly conserved genes across vertebrates in terms of sequence and location and developmental time point of expression [57–60]. Originally disclosed as the first gene identified to be linked to Mendelian speech disorders, it has drawn a great deal of attention as a model to study evolution across vertebrates broadly and related to speech and vocalization in particular [61–69]. While some research has sought out downstream targets of FOXP2 [70–72] or identified its role in developing brain [73–75], lung [76], and other tissues [77–78], there have been fewer reports on regulation of *FOXP2* expression itself [79–80]. This may be partially explained by the complex patterns of regulation and expression: at least four separate TSSs can be active depending on the system under study, and further regulation can occur in *cis* from genomic elements 3 Mbp away from the upstream promoter [81]. Understanding the molecular factors contributing to this complex regulation has relevance to human health as altered FOXP2 expression, both upregulation [82–83] and downregulation [84–86], has been tied to various types of cancer and poor patient prognosis [87]. The high degree of conservation *FOXP2* shows in its transcriptional regulation coupled to its phenotypic relevance in vertebrate development and human health make it an ideal model system to study the principles governing alternative promoter selection.

We used genetically targeted proximity labeling with dCas9-APEX2 to covalently tag the promoter-proximal proteome of the active, upstream promoter (TSS1) of the single-copy locus *FOXP2* in live HEK293 cells with biotin. After streptavidin magnetic bead-based purification with stringent washing and on-bead digestion, we utilized online 2D-LC deep fractionation with TMT-enabled quantitative proteomics via RTS-SPS-MS3 on a tribrid mass spectrometer. We performed bioinformatic analysis on the 6,039 proteins inferred by Proteome Discoverer [88] and identified 373 proteins significantly enriched (Storey-*q* <.05, FC>1.2) at the *FOXP2* promoter compared with the untargeted no gRNA control. This set of proteins was significantly enriched for transcription factors and spliceosome components. To benchmark our candidate proteins and provide a comprehensive characterization of potential regulators of the upstream promoter of *FOXP2*, we compared our proteomic results to multiple orthogonal experimental techniques, including computationally predicted transcription factors recognizing the targeted region and the 222 ENCODE ChIP-Seq experiments performed in HEK293(T) cells (217 transcription factors, 5 histone modifications) [89–92]. We identified and verified several candidate regulators of the chromatin region at the active promoter of *FOXP2* in a discovery-based manner. We also demonstrate that IRF2BP2 negatively regulates *FOXP2* expression.

## Results

### Preparing the FOXP2 Promoter-Proximal Proteome for Mass Spectrometric Analysis by Genetically Targeted Proximity Labeling

The experimental approach is outlined in **Figure 1a**. We began our investigation by generating stable HEK293 cells expressing dox-inducible dCas9-APEX2. Only a small fraction of cells responded to doxycycline treatment after puromycin selection, so we used FACS to generate a monoclonal population to minimize background signal from unlabeled chromatin in non-expressing cells for downstream experiments (**Supporting Figure 1a**). After confirming competence in proximity labeling (**Figure 1b**), we generated three stable cell lines each expressing a different gRNA within the active *FOXP2* promoter region. We performed ChIP-qPCR against the FLAG tag on the proximity labeling construct to confirm accurate targeting using the 2^-ΔΔCt^ approach [93] (See Methods). The no gRNA negative control showed no enrichment after ChIP compared to an off target *GAPDH* control locus (FC=1.15±0.15, where FC=1 is no change) (**Figure 1c**). By contrast, the pooled on-target gRNAs showed significant enrichment after ChIP (FC=2.06±0.31, *p*<.05) (mean±SEM, n=6). This degree of enrichment is consistent with other locus-specific chromatin isolation qPCR results [37,40,41]. We anticipate this to represent a lower bound of enrichment given that streptavidin-based purification enriches on-target loci by up to 284-fold compared with FLAG-based enrichment [35, 94].

**Figure 1.**
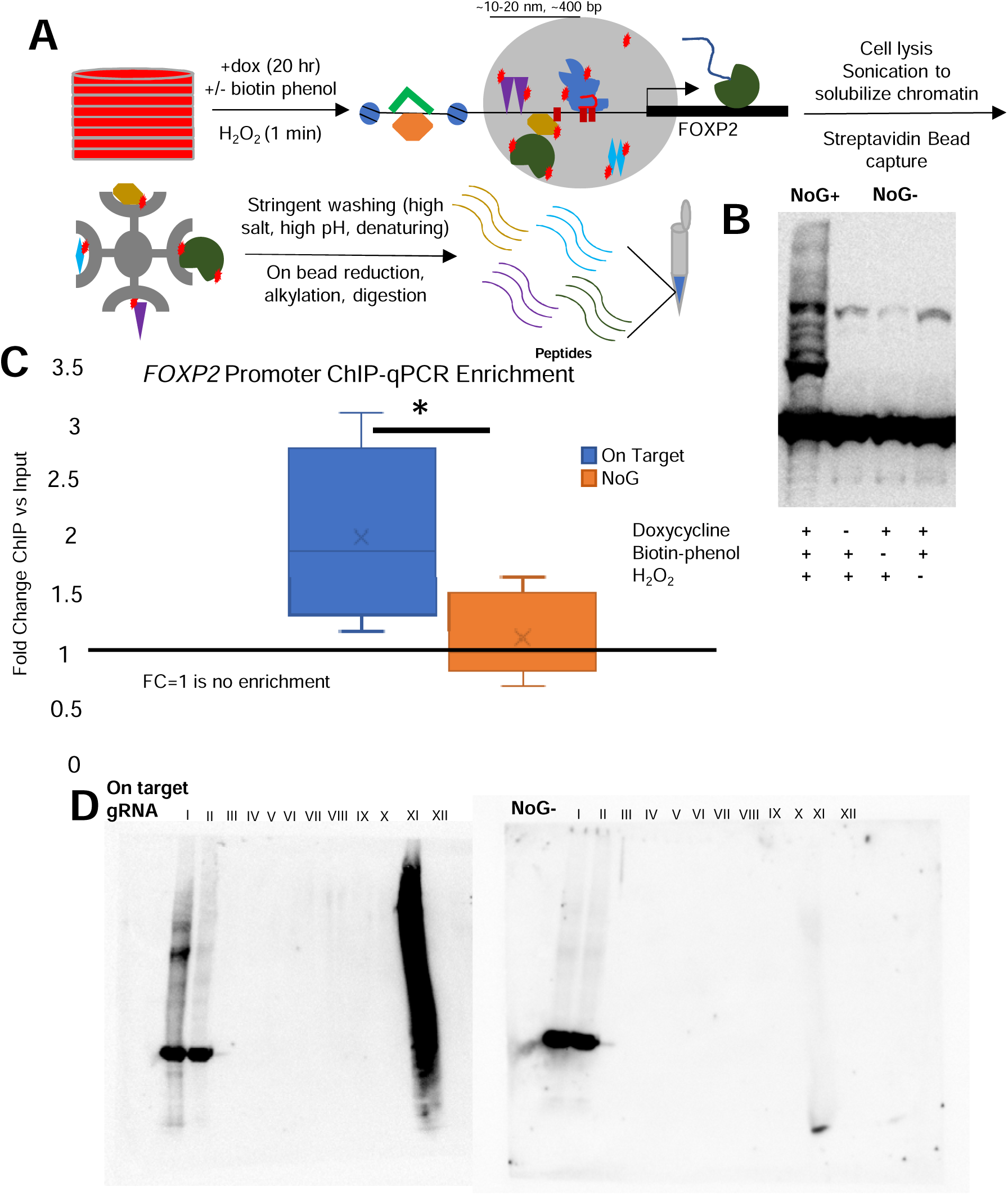
Preparing the *FOXP2* Promoter-Proximal Proteome for Mass Spectrometric Analysis by Genetically Targeted Proximity Labeling. **(a)** Schematic overview of experimental approach for promoter-pulldown proteomics. A large number of input cells (8 15-cm^2^ plates per condition per replicate) are used to generate sufficient material for MS analysis. Activation of genetically targeted proximity labeling with dCas9-APEX2 creates a cloud of reactive phenoxyl radicals (gray circle) that covalently biotinylate proteins (red stars) within 10-20 nm of the promoter targeted by the designed sgRNAs (designated by red boxes). Sonication to solubilize chromatin enables capture of biotinylated proteins by streptavidin magnetic beads. Stringent washing removes background contaminants before on-bead digestion for mass spectrometric analysis of generated peptides. **(b)** Western blot showing proximity labeling positive and negative controls. All samples show the GAPDH loading control and endogenous 75 kDa biotinylated proteins. Systematic exclusion of necessary chemicals prevents proximity labeling, visible as the smear in the condition with all reagents included. The condition with all labeling reagents and no gRNA (NoG+) and the condition with exclusion of biotin phenol (NoG-) are representative of negative controls in the MS experiment. **(c)** Enrichment of targeted *FOXP2* promoter using ChIP-qPCR by the 2^-ΔΔCt^ method. The region targeted by the sgRNAs was enriched by 2.06±0.31, while the untargeted NoG+ condition showed no enrichment (1.15±0.15) compared to an off-target *GAPDH* control. * *p*<0.05, n=6 **(d)** Western blot demonstrating capture and retention of biotinylated proteins during stringent washing for targeted labeling (gRNA2, left) and untargeted no labeling (NoG-, right) conditions. GAPDH loading control is visible in both conditions, while biotin tagging visualized by Streptavidin-HRP is only present in labeling positive conditions. The biotinylated proteins from input are efficiently captured by the streptavidin beads and show no elution during wash steps. 1% of beads pre-digestion demonstrate labeled proteins remain captured, while the absence of signal in the final lane indicates on-bead digestion went to completion. I: input; II: flow-through; III: RIPA wash 1; IV: RIPA wash 2; V: 1M KCl wash; VI: 100 mM Na_2_CO_3_ wash; VII: 6M GdCl wash; VIII: denaturation, reduction, alkylation; IX: TEAB wash 1; X TEAB wash 4; XI: 1% bead elution pre-digestion; XII: remaining bead elution post-digestion.

After confirming successful targeting to the *FOXP2* promoter, we covalently tagged the proximal proteome with biotin using standard proximity labeling conditions. We sonicated chromatin to an average size of ∼450 bp to solubilize labeled chromatin proteins (**Supporting Figure 1b,c**) and captured them with streptavidin magnetic beads. To reduce capture of non-specific background, we increased the stringency of wash steps. Specifically, we replaced the standard 2M urea wash with 6M GdCl. Proteins adopt a random coil conformation at 6M GdCl, whereas the midpoint of unfolding is at 3M urea [95]. Given that the half time for biotin-bound streptavidin unfolding in 6M GdCl is 50 days [96], we hypothesized a wash with 6M GdCl would remove more contaminants while maintaining capture of biotinylated proteins. Furthermore, GdCl can be removed with downstream desalting sample preparation for LC-MS to limit ion suppression in contrast to urea. Accordingly, we observed no loss of material eluting in flow through by Western blot before the final step (**Figure 1d**). We performed on-bead denaturation, reduction, alkylation, and digestion to generate peptides for mass spectrometric analysis.

### Online Deep Fractionation and Quantitative Proteomics via 2D-LC-TMT-RTS-SPS-MS3

The full mass spectrometry workflow is outlined in **Figure 2a**. In addition to the three individual gRNA cell lines tiling the *FOXP2* promoter, we included two negative controls: no gRNA with exclusion of biotin phenol to account for endogenously biotinylated proteins to generate a high confidence contaminant list (NoG-) and a no gRNA condition with biotin phenol to control for background labeling (NoG+) (see **figure 1b**). Each condition was comprised of eight 15-cm^2^ plates and had three independent biological replicates, comprising more than a billion cells (10^9^, although only 60% represent on target chromatin). The fifteen conditions plus a pool of equal amounts of each sample were labeled with TMTs for a 16-plex experiment and 15 μg of protein was injected for LC-MS.

**Figure 2.**
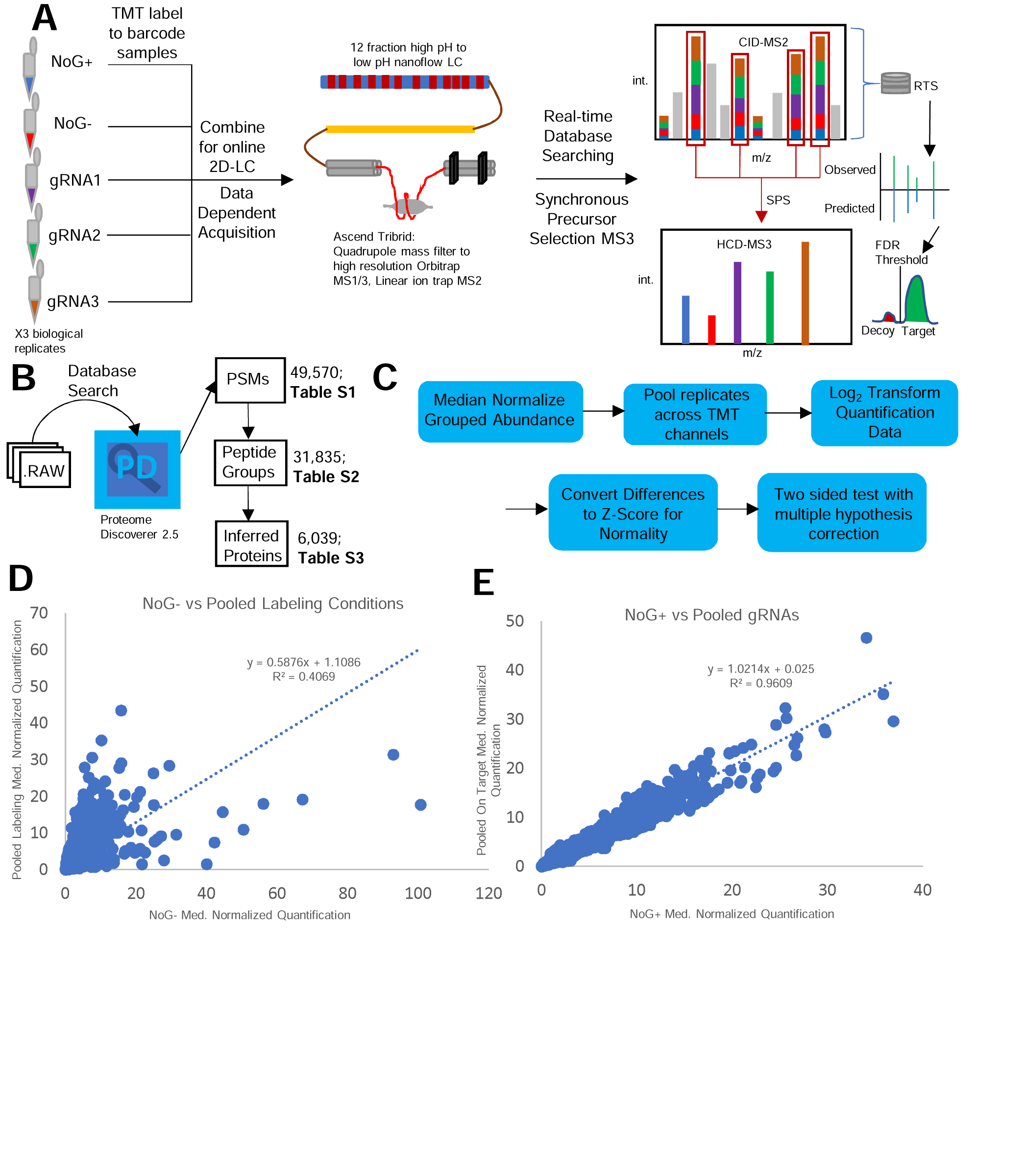
Online Deep Fractionation and Quantitative Proteomics via 2D-LC-TMT-RTS-SPS-MS3. **(a)** verview of 2D-LC-TMT-RTS-SPS-MS3 mass spectrometry workflow. Three biological replicates representing five conditions (three on target sgRNAs, plus two negative controls) are barcoded with TMTs before combination and injection on the LC-MS. A high pH 12-fraction gradient followed by an analytical low pH column separates peptides by retention time before direct connection to the tribrid MS instrument. A quadrupole selects ions in a defined *m/z* range for a high resolution orbitrap scan to capture MS1. Ions are sent to the linear ion trap for MS2 and eventually return to the orbitrap for MS3 quantification. Fragments in MS2 are compared in real-time on the instrument to a protein database to ensure only signals that can be matched to peptides with high confidence are sent to MS3 to economize on the instrument duty cycle, and multiple precursors are selected. Peptide-level FDR is controlled with a decoy database strategy. **(b)** Raw files from the mass spectrometer were processed using a database search via Proteome Discoverer. Total number of identified PSMs (49,570; Table **S1**), peptide groups (31,835; Table **S2**), and inferred proteins (6,039; Table **S3**) are indicated. **(c)** Bioinformatic workflow to analyze protein quantification. **(d)** Global pairwise protein quantification comparing NoG-against all conditions in which labeling occurred (pooled gRNAs + NoG+). There is low correlation in protein quantification (R^2^=0.41) between NoG- and conditions where labeling occurred. **(e)** Global pairwise protein quantification comparing NoG+ against pooled on target conditions. There is a high degree of correlation in protein quantification (R^2^=0.96) between on target and untargeted conditions.

In order to prevent the sample losses that occur on the microscale [97] while maintaining the benefits of deep proteome coverage from extensive sample fractionation, we utilized online 2D-LC directly connected to the mass spectrometer. A 12-fraction high-pH gradient separated peptides before transfer to a low pH nanoflow column (**Supporting Figure 2**). Using RTS-SPS-MS3 for quantitative proteomics (see methods), we detected 49,570 peptide spectral matches (PSMs) (**Supporting Table S1**). We utilized Proteome Discoverer to infer proteins consistent with the 31,835 identified peptide groups (**Supporting Table S2**), generating a list of 6,039 proteins (**Figure 2b**) (**Supporting Table S3**). Since this was a discovery-based effort aimed at low expression proteins (e.g. transcription factors), we did not require at this point that inferred proteins had multiple identified peptides to be included in downstream analysis.

We excluded non-quantified proteins from downstream analysis rather than impute missing values, resulting in a list of 5,074 proteins. We first normalized the grouped abundance of each protein by the median value for each TMT channel [98] and then averaged biological replicates together (**Figure 2c**). Pairwise comparisons of global protein quantification across different conditions revealed the NoG-measurements poorly correlated with all conditions in which labeling occurred, as expected (**Figure 2d**). In contrast, there was a high degree of correlation between the NoG+ conditions and the pooled on target conditions (**Figure 2e**). We considered all proteins that were enriched by at least 1.2-fold in NoG-vs. the pooled labeling conditions and an adjusted *p*-value<.05 to be contaminants and removed them from downstream analysis, resulting in 4,377 proteins. We used Bonferroni correction in generating the contaminant list in order to minimize the number of false positives identified as contaminants while correcting for multiple hypothesis testing [99].

### FOXP2 Promoter Proximal Proteome is Enriched for Transcription Factors and Splicing Machinery

To determine the proteins enriched at the active *FOXP2* promoter, we compared the quantification between the pooled on-target conditions with the untargeted negative control (NoG+). We used the Storey method [100], which is a sharper tool compared to the relative bluntness [101] of the commonly used Benjamini-Hochberg method [102], to correct for multiple hypothesis testing (see methods), generating 596 proteins with *q*<.05. Of those, 373 were enriched with FC>1.2. A recent report utilizing dCas9-APEX2 to identify a promoter-proximal proteome identified biologically relevant proteins that were non-significantly enriched [37]. Ignoring significance, 775 proteins were enriched with FC>1.2. After the filtration steps (**Figure 3a**), we created a volcano plot to visualize the results (**Figure 3b**).

**Figure 3.**
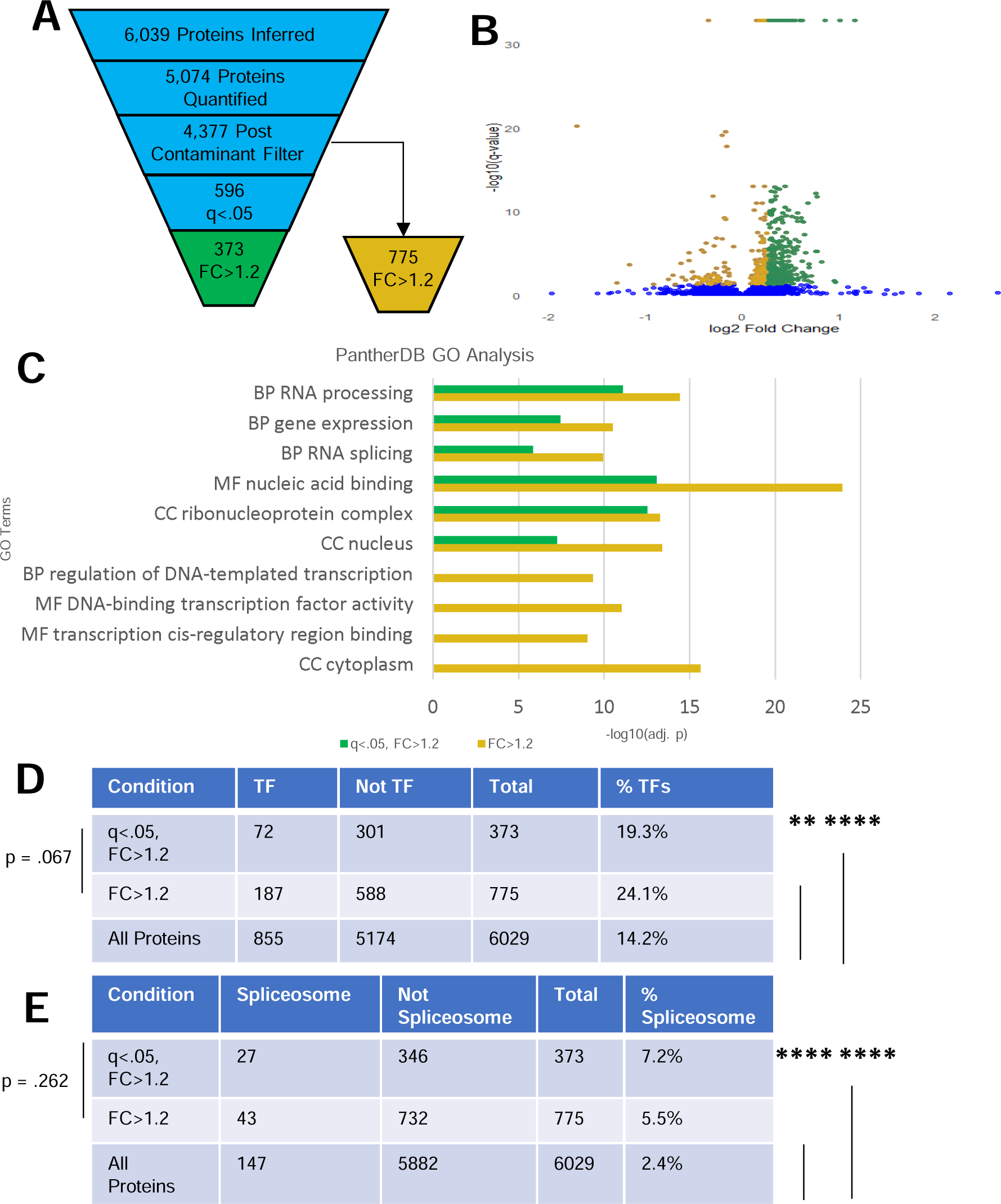
*FOXP2* Promoter Proximal Proteome is Enriched for Transcription Factors and Splicing Machinery. **(a)** Number of proteins remaining after each data analysis filter (quantification, NoG-contaminant list, *q*-value filter, FC>1.2). The number of proteins with FC>1.2 post contaminant filter ignoring the *q*-value filter is also shown. **(b)** Volcano plot showing Log_2_FC(pooled on target vs. NoG+) on the x-axis and −log_10_(*q*-value) on the y-axis. Proteins enriched with FC>1.2 and q<.05 are highlighted in green, while proteins with *q*<.05 and FC<1.2 are golden. **(c)** Gene Ontology enrichment analysis using PantherdbGO for proteins enriched by FC>1.2 with (green) and without (golden) the *q*-value filter. BP: Biological Process; MF: Molecular Function: CC: Cellular Component. **(d)** Contingency table for transcription factors identified in proteins enriched by FC>1.2 with and without the *q*-value filter compared with all identified proteins. \*\**p*<.01 \*\*\*\**p*<.00001. **(e)** Contingency table for components of the spliceosome identified in proteins enriched by FC>1.2 with and without the *q*-value filter compared with all identified proteins. **** *p*<.00001.

We analyzed both sets of enriched proteins (FC>1.2 with or without *q*-value filter) via gene ontology analysis using multiple tools (see methods) [103–106]. Representative results from PantherdbGO are shown in **Figure 3c** and results from all tools with −log_10_(adjusted *p*)>6 are enumerated in **Supporting Table S4**. Expected terms such as nuclear localization and mRNA processing indicate nuclear enrichment was successful. RNA splicing and related terms like ribonucleoprotein complex were enriched as well, potentially indicative of accumulation of dCas9-APEX2 in the nucleolus [39] or co-transcriptional splicing regulation [107]. There tended to be greater statistical significance in terms identified in the list of proteins with FC>1.2 without using the *q*-value filter. That list had greater enrichment of terms related to sequence-specific transcription factor binding and activity of RNA Polymerase II, but it also included false positive terms like cytoplasm. We conclude that analyzing proteins enriched below statistical significance can unveil true promoter proximal proteins but that results must be interpreted with caution and validated by orthogonal approaches.

In order to determine if we were able to successfully enrich transcription factors at the *FOXP2* locus, we compared our enriched proteins to the set of all human transcription factors. We first combined two previously compiled lists of human transcription factors and removed duplicates, generating a set of 2,562 proteins (**Supporting Table S5**) [108–109]. We then generated a contingency table to determine if the fraction of transcription factors in the enriched lists were statistically different from the entire set of proteins inferred in our mass spectrometry experiment (**Figure 3d**). The fraction of transcription factors for the enriched proteins with (*q*<.05, FC>1.2; 19.3%) and without the *q*-value filter (FC>1.2; 24.1%) was significantly different from the total list of proteins (14.2%) by the χ^2^ test. The percent of components of the spliceosome [110] in the enriched lists with (*q*<.05, FC>1.2; 7.3%) and without the *q*-value filter (FC>1.2; 5.5%) was statistically different from the spliceosome components in the whole set of inferred proteins (2.4%) (**Figure 3e**) (**Supporting Table S5**).

Within the set of significantly enriched transcription factors we detected POU3F2, which has been shown to drive reporter gene expression from an intronic *FOXP2* enhancer and has a binding site highly conserved across vertebrates upstream of TSS1 [111–112]. We conclude that genetically targeted proximity labeling is able to successfully detect promoter-proximal proteomes, including classic sequence-specific transcription factors, although identifying statistically significant changes of lowly abundant proteins like transcription factors relative to background still remains a challenge.

### Orthogonal Confirmation of Identified Proteins

Selected proteins enriched by FC>1.2 at TSS1 are indicated in **Table 1**. We turned to orthogonal experimental approaches to benchmark our candidate transcriptional regulators identified by mass spectrometry.

**Table 1.** Selected Mass Spectrometry Enriched Proteins at *FOXP2* TSS1.

| Protein | Class | Notes |
| --- | --- | --- |
| POU3F2 | TF | * ^ Conserved binding site upstream of TSS1 [112] |
| GNL3 | TF | * ^ Member of Wnt/β-catenin pathway [126] |
| NFYC | TF | * ^ # |
| GMEB2 | TF | * ^ # |
| RFX5 | TF | * ^ # |
| HOXD9 | TF | * ^ # |
| ELF1 | TF | * ^ # |
| ETV6 | TF | * ^ |
| ZEB1 | TF | * ^ |
| GATA4 | TF | * ^ |
| FUBP1 | TF | * ^ |
| ZC3H8 | TF | * ^ |
| ATF1 | TF | * ^ # Co-binds with CREB1 |
| CREB1 | TF | ^ # Co-binds with ATF1, binds to *FOXP2* e330 in ChIP-chip analysis [117] |
| YY1 | TF | * ^ # ENCODE ChIP-Seq peak at TSS1 |
| YY2 | TF | * ^ ENCODE ChIP-Seq peak at TSS1 |
| ZFHX2 | TF | * ^ ENCODE ChIP-Seq peak at TSS1 |
| ZNF768 | TF | * ^ ENCODE ChIP-Seq peak at TSS1 |
| ZNF580 | TF | * ^ ENCODE ChIP-Seq peak at TSS1 |
| ZEB1 | TF | * ^ ENCODE ChIP-Seq peak at TSS1 |
| NFATC2IP | TF | * ^ NFATC1 predicted by TFBSPred; NFATC proteins (1 and 2) impact β-cell proliferation through FOXP-dependent mechanism [116] |
| IRF2BP2 | TF | * ^ Partner protein IRF2 predicted by PROMO, IRF2BP1 non-significantly enriched |
| SRSF5 | Spliceosome | * ^ |
| PRKRIP1 | Spliceosome | * ^ |
| SLU7 | Spliceosome | * ^ |
| NOVA2 | Spliceosome | * ^ |
| WBP11 | Spliceosome | * ^ |
| BUD13 | Spliceosome | * ^ |
| DHX16 | Spliceosome | * ^ |
| PRDM9 | Chromatin Remodeler | ^ Validated H3K4Me<sub>2</sub> → H3K4Me<sub>3</sub> activity, consistent with promoter chromatin state |
| GATAD2B | Chromatin Remodeler | * ^ NuRD Complex member |
| MBD2 | Chromatin Remodeler | * ^ NuRD Complex Member |
| MBD3 | Chromatin Remodeler | * ^ NuRD Complex Member |
| MTA3 | Chromatin Remodeler | ^ NuRD Complex Member |

\* $q < .05$ , ^ $FC > 1.2$ , #Predicted by computational TF tool

#### Computationally Predicted Transcription Factor Binding to the *FOXP2* Locus

We first used two separate computational tools that predict potential transcription factor binding sites at user-specified DNA regions, TFBSPred [113] and PROMO [114–115]. TFBSPred predicted potential binding of the mass spectrometry-identified hits NFYA^, NFYC*^, KLF9^, RFX5*^, ATF1*^, CREB1^, GMEB2*^, VEZF1^, CEBPA*, and NR2F2*^ (\**q*<.05, ^FC>1.2). The protein NFATC1 was also predicted by TFBSPred. Both NFATC1 and NFATC2 have been shown to impact β-cell proliferation through a mechanism relying on FOXP proteins from a FOXP1/2/4 triple knockout model [116]. While NFATC1 quantification was non-significant and unchanged (*q*>.05, FC=1.05) and we did not detect NFATC2, the related NFATC2IP*^ was significantly enriched.

The mass spectrometry-identified hits NFIX*, MAZ^, NFYC*^, NFYA^, YY1*^, ATF1*^, CREB1^, HOXD9*^, and ELF1*^ were identified as potential binders by the PROMO web tool. Our mass spectrometry results detected the PROMO-predicted protein IRF2, but it was not enriched. However, related proteins IRF2BP1^ and IRF2BP2*^ were both in enriched via quantitative mass spectrometry. The output of predicted binders from both tools is included as **Supplementary Table S6**. We note that the tools considered all possible human transcription factors across all cell types, including factors not expressed in HEK293 cells. Therefore, we would not expect to have high coverage of the predicted binders.

The known co-regulators ATF1 and CREB1 were identified by both computational tools and our mass spectrometry results. Consistent with these observations, a ChIP-chip analysis of CREB1 in HEK cells showed binding at the *FOXP2* locus [117]. Notably, the sequence in the microarray analysis was included before identification of the active upstream promoter targeted in our experiments. It instead represents a downstream enhancer that has been demonstrated to form an active chromatin loop with TSS1 in HEK293 cells [80] (*vide infra*).

#### ENCODE ChIP-Seq Database

We sought to further validate our mass spectrometry results by identifying true positive proteins that bind to the *FOXP2* promoter in HEK293 cells. The gold standard for identifying protein-DNA interactions in live cells is ChIP-Seq and related approaches. The ENCODE database has a collection of ChIP-Seq experiments targeting a diversity of transcription factors across many cell lines (**Figure 4a**). We analyzed the 222 ChIP-Seq experiments performed in HEK293(T) cells to determine transcription factor and histone post-translational modification status at the *FOXP2* gene. We analyzed binding of factors at the 2,893 bp region of TSS1. A 3C study [80] has demonstrated long range contact between TSS1 and the 7,426 bp region that comprises TSS2, TSS3, and a conserved enhancer 330 kbp away from TSS1, which contains the sequence included in the microarray analysis referenced above; we included that 7.4 kbp regulatory region (hereafter: e330) in our analysis. We also considered the intergenic region upstream of TSS1 between *FOXP2* and *PPP1R3A* and whether there were more than three significant peaks in the *FOXP2* gene body (n>3) to characterize chromatin interactors at the genetic locus in HEK293 cells comprehensively (**Figure 4b**). We compared the presence of significant ChIP-Seq signals at these loci in the ENCODE database with our mass spectrometry data in **Supplementary Table S7**.

**Figure 4.**
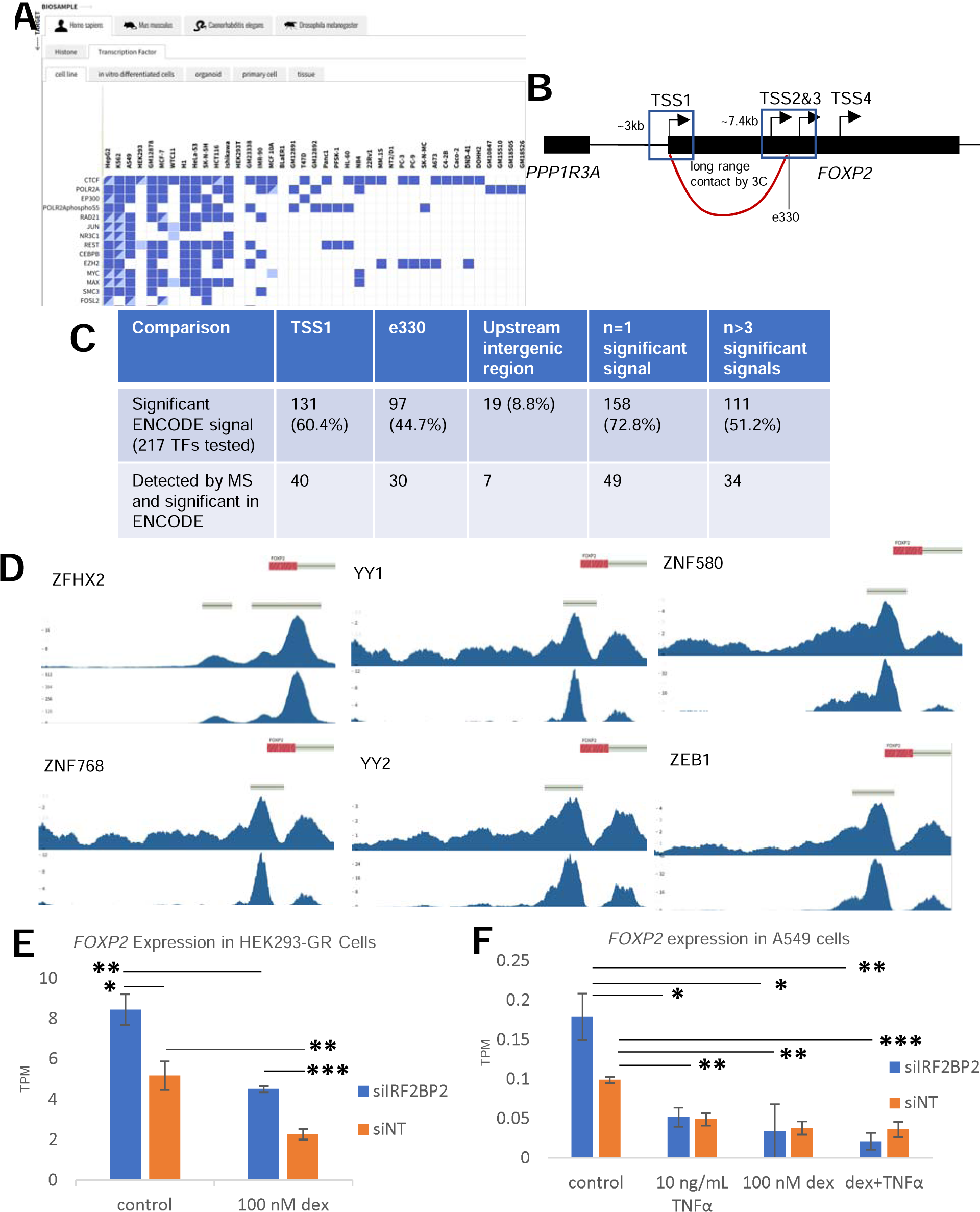
Orthogonal Confirmation of Identified Proteins. **(a)** ENCODE ChIP-Seq Data Matrix available at encodeproject.org. **(b)** Gene body diagram showing *FOXP2* genomic regions examined in ENCODE ChIP-Seq database. TSS1 = hg38 chr7: 114084641-114087534; e330 = hg38 chr7:114411165-114418591, comprising TSS2&3 and a conserved enhancer. A 3C study has indicated these regions form a long range chromatin loop [80], indicated by a red line. **(c)** Number of transcription factors bound at each indicated region of *FOXP2* from ENCODE ChIP-Seq database and corresponding mass spectrometry results. See also Table **S7** for detailed comparison of ENCODE results to mass spectrometry results. **(d)** ENCODE ChIP-Seq tracks of the 6 proteins with called peaks that were enriched in the mass spectrometry dataset (*q*<.05, FC>1.2). Each track shows the *FOXP2* gene body, the IDR (Irreproducible Discovery Rate) called peak thresholds, the fold change of the indicated transcription factor over control, and the associated *p*-value. Each transcription factor has its own data-dependent scale bar for fold-change and *p*-value. **(e)** Expression of *FOXP2* in transcripts per million (TPM) in HEK293 cells constitutively expressing the glucocorticoid receptor (GR). Expression of *FOXP2* transcripts are increased upon siRNA knockdown of IRF2BP2 and decreased upon activation of GR signaling by dexamethasone. Error bars = SEM, n=4. * p<.05, ** *p*<.01, *** *p*<.001. **(f)** Expression of *FOXP2* in TPM in A549 cells before and after siRNA knockdown of IRF2BP2 or activation of GR and/or TNFα signaling. Error bars = SEM, n=3. \**p*<.05, \*\**p*<.01, \*\*\**p*<.001.

Our targeted promoter showed H3K4Me_3_ and H3K27Ac histone marks, consistent with an active TSS [118]. These histone modifications were present at e330 in addition to H3K4Me_1_, in line with an active enhancer region. Consistent with the observed chromatin state of our targeted promoter, we detected the non-significantly enriched chromatin remodeler PRDM9^, which has validated H3K4Me_2_ ➔ H3K4Me_3_ methyltransferase activity [119]. We also detected members of the Nucleosome Remodeling and histone Deacetylation (NuRD) complex [120] GATAD2B*^, MBD2*^, MBD3*^, and MTA3^. By contrast, we failed to detect enrichment of multiple members of the SWI/SNF, ISWI, and INO80 families of chromatin remodelers, suggesting the NuRD complex may regulate the epigenetic state of the *FOXP2* promoter in HEK293 cells.

Out of the 217 transcription factors studied in HEK cells by ENCODE, 131 (60.4%) showed significant ChIP-Seq signals at TSS1 (**Figure 4c**). At e330, 97 (44.7%) transcription factors showed a significant signal. Only 19 (8.8%) showed binding in the upstream intergenic region between TSS1 and *PPP1R3A*. There were 158 factors (72.8%) with any significant signals in the *FOXP2* gene body, while only 111 (51.2%) had more than three significant signals. Most of the transcription factors detected at e330 overlapped with those found at TSS1, with the exception of 12 proteins that had no promoter peak but were bound to the enhancer (FOXA1, TRIM28, ZNF384, MEIS1, ZFP3, ZNF34, ZNF362, ZNF624, AEBP2, GFI1B, GLI2, ZNF654). Out of the 217 transcription factors that have ChIP-Seq experiments in HEK293 cells in the ENCODE database, we detected 71 by mass spectrometry. We only detected 40 of the transcription factors with ChIP-Seq signals at TSS1 in our mass spectrometry dataset, meaning 91 (69.5%) of ChIP-Seq positive transcription factors in the ENCODE database were undetected as false negatives. On the other hand, 56% of the mass spectrometry detected transcription factors with ENCODE data (40/71) were true positives before performing bioinformatic analysis on the dataset.

Out of the 71 ENCODE transcription factors detected in our data set of 6,039 inferred proteins, only 57 were enriched in labeled conditions vs. NoG-; the majority of proteins eliminated (12/14, 86%) were not quantified at all, indicating they were present near the detection limit of the instrument rather than being enriched in NoG-. There were 13 transcription factors with *q*<.05, of which 11 had FC>1.2. Only 8 of those 13 (61.5%) had ChIP-Seq signals, and the two eliminated by the FC>1.2 filter (TARDBP and PKNOX1) were true positives in ENCODE ChIP-Seq. Out of these false positives, one (ZNF384) had a signal at e330, while 3 had weak underlying mass spec data (e.g. degenerate peptides [peptide(s) could derive from multiple proteins, i.e. not unique] or identified by a single peptide, “one hit wonders”) and could be reasonably discarded, leaving one false positive (7.7%). The ChIP-Seq tracks of the 6 proteins with *q*<.05, FC>1.2, and significant ENCODE ChIP-Seq signals are shown in **Figure 4d**. Comparing ENCODE results to the list of all proteins with FC>1.2 without the *q*-filter, 25 of 57 were enriched, but only 14 showed signals at TSS1 (56%). Out of the 11 false positives, there were 3 transcription factors with peaks at e330 (TRIM28, ZNF384, and MEIS1), and 6 had weak underlying mass spectrometry data. Given the confirmed chromatin loop between e330 and TSS1 via 3C studies [80] and the proximity-based nature of our labeling method, we posit that some of the identified e330 binders may be true positives rather than false positives.

#### Comparison to Known Regulators of *FOXP2* Expression

We further compared our mass spectrometry data and results from the ENCODE database with the few known regulators of *FOXP2* expression. One of the most well-known and best characterized regulators of *FOXP2* expression is the Wnt/β-catenin transcription factor LEF1. Experiments in zebrafish embryos have demonstrated that lef1 directly binds to and regulates the expression of *foxp2* [121]. In contrast, LEF1 was detected but not enriched in our mass spectrometry results, and ENCODE showed no significant ChIP-Seq signal at TSS1 or e330 in HEK293 cells. This observation highlights the need for care in transferring observations of transcriptional regulation across cell types. On the other hand, the TCF/LEF transcription factor family member TCF7L2 showed a significant ChIP-Seq signal at TSS1 and e330. Although we detected TCF7L2 in our inferred proteins, it was not significant and was more enriched in NoG+ conditions compared to on target. However, analysis of the PSMs showed that the only identified peptide was shared with LEF1 (degenerate “one hit wonder”). This observation highlights the difficulty in the protein inference problem for which there are multiple approaches and no commonly accepted best practice [122–125]. It is noteworthy that one of the top hits in our list (*q*<.05, top 10 FC vs NoG+), GNL3, is a component of the Wnt/β-catenin signaling pathway [126].

The transcription factor FOXK2 was predicted to bind to the *FOXP2* promoter by TFBSPred and showed a significant signal in the ENCODE ChIP-Seq database at TSS1. However, there was no difference between on and off-target conditions in our quantitative dataset (FC=1.02). Furthermore, the protein ZBTB20 showed a significant ChIP-Seq signal, and the mouse homolog Zbtb20 has been shown to bind to and control *FoxP2* expression in the developing mouse brain [127], but we failed to detect it in our mass spectrometry data set. The homolog of the human protein PAX6 has been identified as binding to the *foxp2* locus and impacting gene expression in zebrafish models [128]. We detected this protein but it was not enriched in on target conditions (FC=1.05), although it was not tested in HEK293 cells by ENCODE. Each of these observations highlights that a lack of detection or enrichment in our mass spectrometry data set does not preclude a protein from being a member of the *FOXP2* promoter-proximal proteome.

#### IRF2BP2 is a Negative Regulator of *FOXP2* Transcription

Given that there can be limited overlap between transcription factor binding and regulatory activity [129], we sought to demonstrate a direct impact on transcriptional output rather than simply a binding interaction at the *FOXP2* promoter from within our proposed candidate regulators. The protein IRF2BP2 was significantly enriched in our proteomic assay and has been identified as an IRF2-dependent transcriptional repressor [130–131]. Given that IRF2BP1 was also enriched (non-significantly) and IRF2 was predicted from the PROMO binding tool, we sought to characterize effects of IRF2BP2 on *FOXP2* expression. Characterization of IRF2BP2 genome-wide binding and effects of its depletion on transcription have been reported previously in HEK cells that constitutively express the glucocorticoid receptor [132].

As shown in **Figure 4e**, siRNA knockdown of IRF2BP2 leads to a significant 1.6-fold increase in expression of *FOXP2* compared with a non-targeting control (siNT). This effect persisted when glucocorticoid signaling was activated by addition of 100 nM dexamethasone. Notably, activation of glucocorticoid signaling by dexamethasone significantly decreased *FOXP2* expression in both siIRF2BP2 and siNT conditions. There were three called peaks in the *FOXP2* gene body for IRF2BP2 binding via ChIP-Seq in control conditions and only one peak with dexamethasone treatment, although the control peaks sat between TSS1 and e330 and did not include our targeted promoter. Conspicuously, however, one of the three peaks in control conditions sat approximately 56 kbp downstream of the targeted promoter, a region that has been shown to form a weak chromatin loop with TSS1 across neuronal cell types [80]. The peak in dexamethasone-treated cells was more than 400 kbp downstream of TSS1 and did not have evidence for long range chromatin looping in any cell types since it was outside the interrogated enhancer-promoter pairs in the 3C study.

To determine whether these observations were translatable to other cellular contexts, we compared *FOXP2* expression before and after knockdown of IRF2BP2 in the lung cancer cell line A549 from the same study (**Figure 4f**). FOXP2 is only weakly expressed at the transcript and protein level in A549 cells, lowering power to call statistical significance (note scale bar differences between **4e** and **4f**). This downregulation is tied to increased aggressiveness of lung cancer [133]. Knockdown of IRF2BP2 led to a 1.8-fold increase in *FOXP2* expression that approached but did not reach significance at the *p*<.05 level compared to siNT conditions (p=.0558). Treatment with dexamethasone, TNFα, or a combination of both significantly decreased *FOXP2* expression. These changes did not differ between siIRF2BP2 and siNT conditions (p>.3), and there were no detectable differences among the individual or combined treatments. We conclude that IRF2BP2 can negatively regulate *FOXP2* expression across cell types, as can glucocorticoid and TNFα signaling.

## Discussion

The presence of multiple promoters within a gene permits regulatory systems that can incorporate complex logic operations for precise titration of gene dosage [134]. While synthetic biologists are just beginning to harness these principles for designed cellular phenotypes, evolution has relentlessly optimized gene circuits as long as they have existed [135]. Ultraconserved elements with greater than 90% sequence identity for more than 100bp emerged more than 400 million years ago, before the evolution of lobe-finned fish and amphibians [136]. These are enriched in non-coding regulatory regions, and *FOXP2* harbors a cluster of more than 10 highly conserved regions shared across vertebrates but not invertebrates [137]. This high conservation allows FOXP2 to fill many roles, from neuronal expression related to vocal learning [69] to jaw formation and body plan development [76, 138].

In an effort to continue to unravel this highly conserved regulatory module, we sought to deeply characterize in HEK293 cells the proteins interacting with the upstream promoter active in that and many other cell types [79] using genetically targeted proximity labeling via dCas9-APEX2. The extensive washing to remove background contaminants coupled with quantitative mass spectrometry via online 2D-LC-TMT-RTS-SPS-MS3 enabled by a tribrid mass spectrometer allowed us to attain deep proteome coverage and identify 373 proteins significantly enriched at the *FOXP2* promoter (Storey-*q*<.05, FC>1.2), including 72 transcription factors. Many of these candidate transcriptional regulators had orthogonal evidence of biological relevance – from previous literature evidence to computationally predicted transcription factor binding to ENCODE ChIP-Seq data. A great deal of effort has gone into generating mice with disruptions to FoxP2 expression in a bid to understand its biological function [139–142]. The candidate regulators identified in this study can be used for hypothesis generation to uncover the upstream mediators of FoxP2 effects in different conditions and developmental time points.

All identified proteins are only candidate regulators until confirmed through orthogonal lines of evidence. We leveraged the ENCODE database to provide direct evidence for or against factors that were present in both data sets. Although there were more than 200 transcription factor ChIP-Seq experiments in HEK293 cells, some of our candidate regulators that were not tested in HEK cells had been tested in other cell lines. Future locus-specific chromatin isolation studies would benefit from utilizing one of the ENCODE ‘depth’ cell lines, like HepG2 or K562, if they accurately recapitulate the transcriptional conditions of interest to researchers.

Confirmation of candidate regulators via ChIP-Seq or ChIP-qPCR is a resource and labor intensive process that also requires the existence of ChIP-grade antibodies. Contemporary locus-specific chromatin studies often follow up on only a few biologically informed hits and deeply characterize them [26, 36, 37, 41]. A fruitful frontier is determining changes in proteome composition at a targeted locus in response to experimental perturbations [38–40]. Mass spectrometry provides rich, multidimensional data of measured peptides and their quantification to the protein level which can be used to prioritize hits for follow up in a discovery-based manner. Some of the false positives identified in our mass spectrometry dataset that did not show signals in the ENCODE database had weak underlying PSM data (identified by degenerate peptides and/or “one hit wonders”), highlighting the utility of incorporating the mass spectrometry data at all levels for decision making. There is much to be gained from systematic analysis of proteomics datasets in addition to targeted follow ups. Caution must be taken, however, especially with results from labs without deep MS-based proteomics experience – it is not uncommon in the proteomics field to see uncorrected *p* values at the protein quantification level [101] despite stringent, data-driven FDR at the PSM, peptide, and protein identification levels. With calls from active practitioners [49, 99] and options for multiple hypothesis correction in most MS data analysis tools, the proteomics community is beginning to adopt statistical strategies for multiple hypothesis testing in protein bioinformatic analysis, but it is by no means universal.

It is noteworthy that LEF1 showed no signal at TSS1 or e330 in ChIP-Seq given its demonstrated role in regulating *Foxp2* expression in developing zebrafish [121]. This was not due to lack of expression – we detected LEF1 and it was enriched relative to NoG-conditions. Nuclear expression of a transcription factor coupled to open chromatin at an actively transcribed gene that it is known to regulate under certain conditions is not sufficient to induce binding. It is possible that LEF1 can be recruited to the *FOXP2* locus in HEK293 cells under the appropriate signaling conditions. Alternatively, LEF1 may need co-regulators to bind to *FOXP2* that are not expressed in HEK293 cells. Another possibility is that LEF1 binding and regulation of *FOXP2* occurs at genomic elements other than the upstream transcribed promoter or active enhancer 330 kb downstream, or that LEF1 regulates *FOXP2* expression without a direct binding interaction [129]. Other components of the Wnt/β-catenin signaling pathway do have evidence for interaction with *FOXP2* TSS1 in HEK cells – TCF7L2 binding at TSS1 was significant according to ENCODE ChIP-Seq, and GNL3 was one of our top enriched hits.

### Limitations of the Study

One limitation of the present study is the use of an untargeted dCas9 as a negative control rather than a non-targeting gRNA containing a sequence not found in the human genome. The dynamics of dCas9 interrogating the genome differs in the presence and absence of a gRNA [143]. An off-target locus [33] or an alternative locus of interest for comparative studies [38] can also be used as a control to account for the high background in proximity labeling studies. Despite the high background, proximity labeling is a valuable approach that provides complementary information to affinity purification, including the ability to detect lower abundance proteins [144]. A snapshot of chromatin regulators within an approximately 400 bp radius [35] over the course of a minute helps unravel the dynamic processes underlying gene regulation.

Our on-bead digestion protocol leaves behind the residues and associated proteolytic fragments that were biotinylated in our sample preparation (canonically tyrosine, but cysteine and lysine labeling have been observed as well [145]), compounding the protein inference problem by removing potentially informative peptides from what we injected on the mass spectrometer. Using desthiobiotin [24, 146] or a clickable biotin analogue with a cleavable linker to release labeled proteins from the streptavidin beads before digestion could capture those peptides while preventing the unacceptable sample loss for low-input studies that occurs with repeated manipulations on the microscale. Such an approach could be adapted in principle to top-down proteomics protocols to study alternative proteoforms interacting with a promoter [147].

Another shortcoming is that this is necessarily a bulk measurement averaged across many cells. It is unlikely that all the identified regulators co-occupy the *FOXP2* promoter at the same time in the same cell, obscuring the important biological insights that can be obtained at the single cell level [148]. Understanding which factors co-bind or which are mutually exclusive to each other is not possible with this data set alone. Split enzymes or FRET assays could be an approach to test co-binding of specific factors.

Another issue is that cells were not synchronized. The transcription factor YY1, significantly enriched in our data set, predicted by PROMO, and with a significant ChIP-Seq signal at TSS1 according to ENCODE (**Figure 4d**), has been shown to have differing effects on its role in establishing and maintaining chromatin loops based on the cell cycle at some genomic locations [149]. The interaction of FOXP2 itself with DNA is also tightly regulated and controlled throughout the cell cycle [150]. Synchronizing cells could help ensure promoters of interest have similar chromatin states.

There were many false negatives (nearly 70% of ENCODE-positive transcription factors) despite the high initial input of cells (∼6*10^8^ on-target chromatin equivalents). Some transcription factors we detected were not quantified, indicating they were present below the limit of quantification. Utilizing mass spectrometry approaches like parallel reaction monitoring could help identify candidate regulators at an isolated genomic locus in a hypothesis-driven, targeted approach [151]. This approach could also be adapted to try to increase sequence coverage from proteins with weak PSM support by prioritizing for MS analysis unique PSMs to solve the peptide degeneracy problem or to determine if “one hit wonders” are capable of generating any further hits.

It is well appreciated that transcription factors can undergo extensive posttranslational modifications that impact subcellular localization, interaction partners, and regulatory function [117, 150, 152–153]. We used a closed search strategy and did not specify functionally relevant posttranslational modifications like phosphorylation in analyzing our data. An open search, spectrum-centric strategy can identify and localize posttranslational modifications even if they have not previously been identified [154–156]. Such an approach could in principle identify peptides and proteins within this dataset containing posttranslational modifications related to transcriptional regulatory function.

Generating enough starting material for deep proteome coverage is labor intensive. Contemporary locus-specific proteomics experiments often target repetitive regions like telomeres (92 copies of repetitive sequence per cell) [24, 25, 31, 33, 34, 39], centromeres (46 copies of repetitive sequence per cell) [31, 34, 39, 157], or LINE-1 transcribed promoters (80-100 copies per cell) [157–158]. Another approach is to use reporter plasmids present at a higher copy number to increase intrinsic signal [37] or to use a model system with a smaller genome to reduce background [159], examples of single-copy locus enrichment (two copies per cell) in human systems notwithstanding [26, 27, 35, 36, 39–41]. Strategies to reduce necessary input volume are needed to enable more widespread adoption of promoter-pulldown proteomics. Single cell proteomics approaches have utilized a TMT-based signal boosting approach to generate enough material for analysis [160]. Adaptation of that paradigm for promoter-proximal proteomes is a promising avenue for development. The data dependent acquisition approach used in this study samples only a fraction of the ions that elute from the LC for MS analysis stochastically. Data independent acquisition approaches, especially when coupled with parallel accumulation-serial fragmentation (dia-PASEF) enabled by ion mobility separation, can sample all ions eluting off the LC and provide deep proteome coverage on material-limited samples, giving another potential avenue to reduce sample input requirements [157, 161–162]. For promoter occupancy studies on very highly conserved genes, Epi-Decoder provides an interesting proteomics-by-sequencing approach that can be used in yeast [163].

## Conclusion

The majority of disease-risk variants identified in genome wide association studies map to non-coding regions [164]. Understanding how these genetic variants give rise to different phenotypes and impact human health requires being able to determine chromatin associated proteins and how they differ across sequence diversity. Forward genetic studies relying on targeted antibodies to determine genome-wide association of disease relevant factors have been leveraged to great effect. Even with increasingly sophisticated protocols that allow individual labs to collect data at a scale previously accessible only to multi-institution consortia [165], these forward genetic studies are intrinsically limited by the availability of high-quality antibodies. Genetically targeted proximity labeling coupled to quantitative mass spectrometry proteomics is a powerful approach for discovery-based reverse genetics to determine the phenotypic impact of sequence variation without the need for ChIP-grade antibodies or a prior hypothesis of molecular mediators. The biochemical and analytical challenges can also serve as a Muse for development of new and improved proteomics instrumentation, data acquisition, and analysis protocols [166–167].

---

**Figure SF1.**
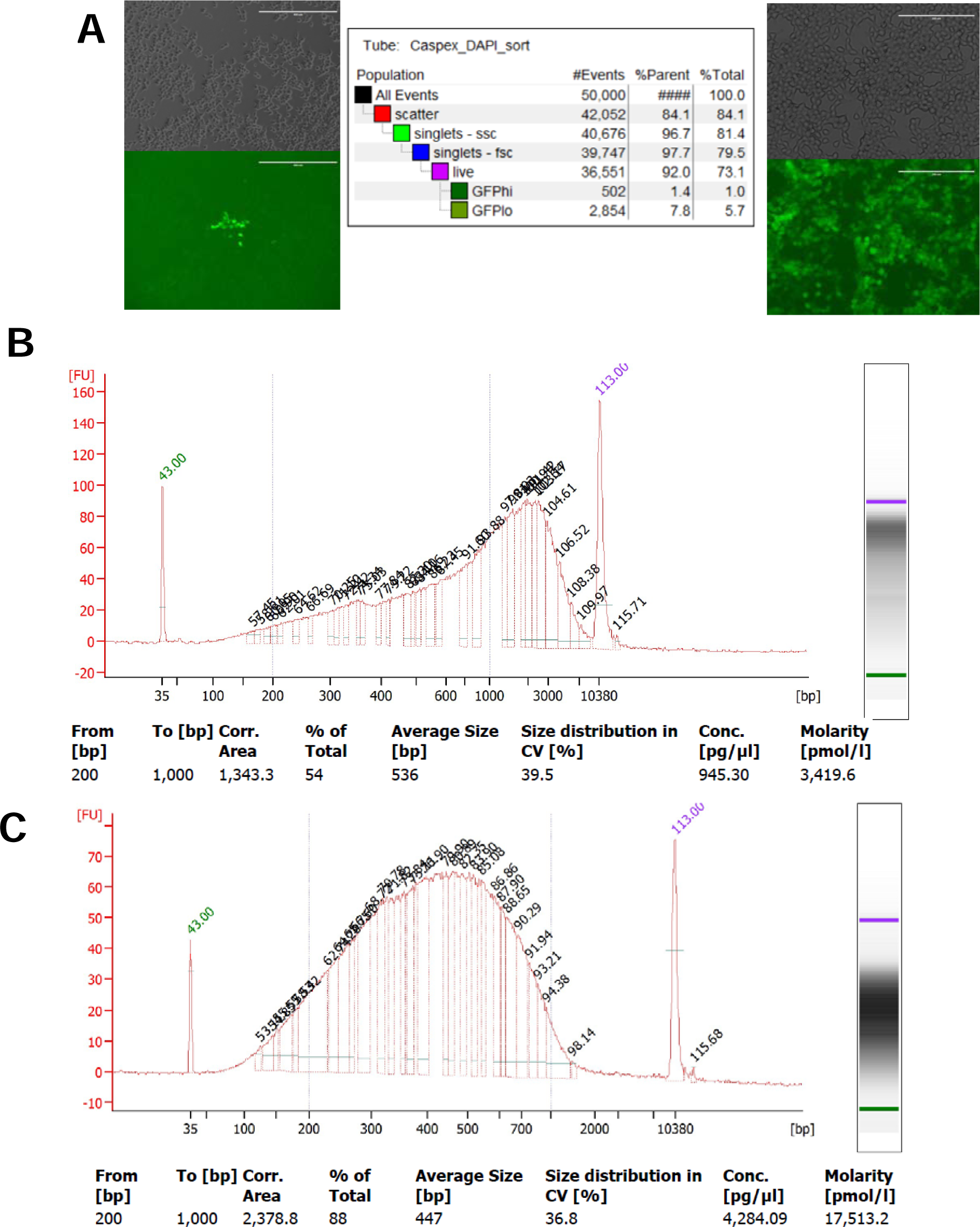
Preparing Chromatin for Analysis. **(a)** Bright field and GFP fluorescence response to 21 hr doxycycline treatment before (left, scale bar 400 μm) and after (right, scale bar 200 μm) monoclonal isolation by FACS. Approximately 5% of polyclonal cells responded to doxycycline treatment before FACS as measured by GFP fluorescence, indicated by the FACS report (center). **(b)** Representative chromatin size distribution by Bioanalyzer analysis after sonication for cross-linked chromatin for ChIP-qPCR. **(c)** Representative chromatin size distribution by Bioanalyzer analysis after sonication for non-cross-linked chromatin for proteomic analysis.

**Figure SF2.**
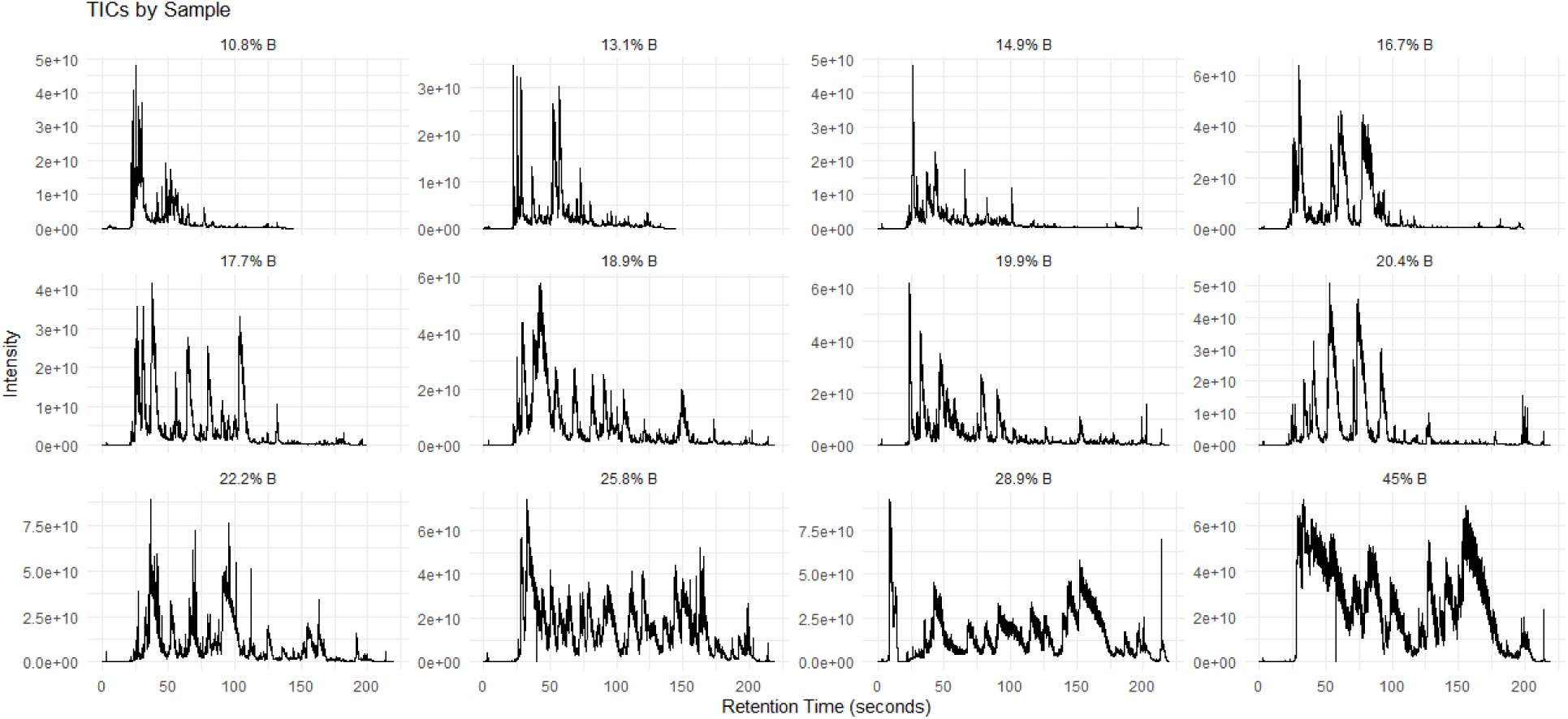
Total Ion Chromatograms from Each High pH Fraction from 2D-LC. TICs produced with RaMS package in R from the mzML file corresponding to each high pH fraction.

## Acknowledgements

This work was performed on the ancestral land of the Muwekma Ohlone Tribe. Cell sorting for this project was done on instruments in the Stanford Shared FACS Facility (RRID: SCR_017788). The Stanford University Department of Statistics Consulting Services and the Vincent Coates Foundation Stanford University Mass Spectrometry facility (RRID:SCR_017801) provided helpful feedback. The authors would like to acknowledge members of the Snyder lab at Porter Drive, particularly Y Zhu, S White, and M Nshanian, for helpful discussions. The authors appreciate comments on early drafts provided by S Saad, S Nageshwaran, and G Erwin.

## Author Contributions

Conceptualization – TMGM; Data Curation – TMGM; Formal Analysis – TMGM; Funding acquisition – MPS; Investigation – TMGM, LR, and RJ; Methodology – TMGM and LJ; Project Administration – TMGM, LR, and LJ; Resources – MPS; Supervision – MPS; Visualization – TMGM; Writing – Original Draft Preparation – TMGM; Writing – Review and Editing – all authors.

## Method Details

### Cell Line Construction and Culture

HEK cells were cultured in Dulbecco’s Modified Eagle Medium containing 10% FBS and 1% penicillin-streptomycin and passaged every 3 days. Cells were maintained at 37 °C in a humidified atmosphere containing 5% CO_2_. To stably incorporate the Caspex construct, cells were transfected with Lipofectamine 3000 (Invitrogen) according to the manufacturer’s instructions and selected for two weeks with 4μg/mL puromycin in the media. Inducible Caspex expression was a gift from Steven Carr & Samuel Myers (Addgene plasmid #97421) [35]. Single colonies were selected by FACS at the Stanford Shared FACS Facility and tested for doxycycline inducibility of Caspex monitored by GFP visualization and anti-FLAG Western blotting with Rabbit M2 (Sigma F2555). Individual sgRNA constructs were transfected into the best responding Caspex cell line and selected with 200 μg/mL hygromycin. Cell lines were maintained with 4 μg/mL puromycin and/or 200 μg/mL hygromycin to maintain selection pressure for the Caspex and sgRNA plasmids, respectively, until use in ChIP-PCR or proximity labeling experiments.

### Plasmid Construction

The UCSC Genome browser was utilized to select sgRNA sequences targeting the promoter of *FOXP2* that conformed to requirements for expression from the U6 promoter while minimizing off target effects. Selected guides were required to start with G for transcription initiation and were rejected if they contained a run of 4 or more T’s in a row to prevent early termination. The MIT specificity score for all guides was greater than 90. Appropriate gRNA sequences were cloned into the lenti-sgRNA hygro backbone, a gift from Brett Stringer (Addgene plasmid #104991) [168]. Briefly, the lenti-sgRNA hygro plasmid was digested with *BsmBI* (New England Biolabs) to create overhangs that could hybridize with Fwd-5’-ACACCGN_20_G-3’ and Rev-5’-AAAACN_20_G-3’ sequences, which were ligated with the designed oligonucleotides using T4 polymerase (New England Biolabs). The vector was transformed into One Shot Stbl3 chemically competent *E. coli* (Invitrogen), single colonies were picked, and plasmid DNA was purified with a QIAGEN EndoFree Maxiprep Kit. Successful gRNA incorporation was confirmed by Sanger sequencing (Integrated DNA Technologies). Primers for gRNA incorporation were synthesized by Integrated DNA Technologies and were as follows: g1 Fwd-5’-ACACCGCAGACACCTTTCGGTGATAG-3’, Rev-5’-AAAACTATCACCGAAAGGTGTCTGCG-3’; g2 Fwd-5’-ACACCGACACCTTTCGGTGATAGGGG-3’, Rev-5’-AAAACCCCTATCACCGAAAGGTGTCG-3’; g3 Fwd-5’-ACACCGTTATCCCGAAGCGTCAGTAG-3’, Rev-5’-AAAACTACTGACGCTTCGGGATAACG-3’ to target sequences in chr7 (hg38) of gRNA1:114087491-114087513, gRNA2:114087494-114087516, gRNA3:114087279-114087301. Target sequences were chosen for proximity to the transcription start site and overlapping labeling radii (∼400 bp [35]) within the *FOXP2* TSS1 promoter while conforming to U6 expression requirements.

### Cross-Linking and Sonication for ChIP

Doxycycline (70% in ethanol) was added to cells at a final concentration of 1 μg/mL in a 15-cm^2^ plate for 21 hr so Caspex expression would be induced when cells were approximately 90% confluent (∼10^7^ cells). Caspex expression was visually confirmed by fluorescence of the GFP marker. A single cell suspension was crosslinked with 1% formaldehyde at room temperature with rotation for 10 min. To quench formaldehyde, 2M glycine was added to a final concentration of 125 mM and incubated for 5 min at room temperature with rotation. Cells were washed twice with ice cold PBS, pelleted, snap-frozen in liquid nitrogen, and stored at −80°C until use. Cells were thawed at 4°C in ice cold PBS with rotation for 30 minutes. Pelleted cells were treated with 3 mL hypotonic buffer (20 mM HEPES pH 7.9, 10 mM KCl, 1mM EDTA pH 8.0, 10% glycerol) with protease inhibitors (Roche cOmplete protease inhibitor [1 tablet/50 mL], 0.5 mM PMSF) and 1 mM DTT added just before use. The plasma membrane was allowed to swell for 10 minutes before shearing with 30 strokes of a Dounce homogenizer. Total time for swelling and homogenization was limited to 15 minutes. Nuclei were pelleted and washed once with ice cold hypotonic buffer before lysis in 3 mL ice cold RIPA buffer (150 mM NaCl, 50 mM Tris-HCl pH 8.0, 1% IGEPAL CA-630, 0.5% sodium deoxycholate, 0.1% SDS) with protease inhibitors, DTT, and phosphatase inhibitors (1 mM Na_2_P_2_O_4_, 2 mM Na_3_VO_4_, 10 mM NaF) added just before use. Lysed nuclear pellets were sonicated on ice to shear chromatin for solubilization with a SFX250 Sonifier (Branson) set to an intensity (output control) of 3.5. Lysates were sonicated for 16 rounds of 30 s on and 30 s off, taking care to prevent foaming. To prevent sample overheating, lysate was allowed to rest on ice for at least 2 minutes every four cycles. Lysate was clarified by centrifugation at 14,000 rpm for 15 minutes at 4°C. Supernatant was transferred to a 15 mL Falcon tube, diluted to 4 mL total in RIPA buffer, flash frozen in liquid nitrogen, and stored at −80°C until use. Chromatin fragmentation was assessed after de-crosslinking (*vide infra*) using a Bioanalyzer (Agilent). Typical results produced an average size of ∼530 bp with 55-60% of fragments in the 200-1000bp range (**SF1b**).

### Chromatin Immunoprecipitation

For chromatin immunoprecipitation, each biological replicate produced two technical replicates for pulldown. For one technical ChIP replicate, 2 mL of sonicated lysate was used. Before pulldown, 20 μL input (1%) was removed to compare nucleic acid enrichment before and after ChIP. Sonicated lysate was incubated at 4°C overnight with 5 μg FLAG Rabbit M2 monoclonal antibody (Sigma F2555) with rotation. A 1:1 mixture of 150 μL total Protein A:Protein G magnetic beads (Invitrogen Dynabeads, 30 mg/mL each) was washed twice with 1 mL ice cold RIPA buffer to pre-clear the beads. The beads were transferred fully to the antibody-complex-chromatin mixture and incubated for 1 hr at 4°C with rotation. The beads were washed 3 times with ice cold RIPA buffer with inhibitors added followed by a wash with ice cold PBS. The beads were transferred from the 15 mL Falcon tube to a 2 mL DNA lo-bind Eppendorf tube with 800 μL + 2 times 400 μL ice cold PBS to ensure complete transfer. The PBS was removed and beads were incubated with 100 μL of freshly made 1% SDS, 10 mM Tris pH 8.0, 1 mM EDTA at 65°C for 10 minutes with gentle mixing by vortex every 2 minutes. The supernatant was collected and beads were incubated with 150 μL of 0.67% SDS, 10 mM Tris pH 8.0, 1 mM EDTA at 65°C for 10 minutes with gentle mixing every 2 minutes before combining both eluates. Input DNA (stored at 4°C overnight) was diluted 1.5x in 1% SDS, 10 mM Tris pH 8.0, 1 mM EDTA before both ChIP DNA and input DNA had crosslinks reversed overnight at 65°C. An equal volume of 1% SDS, 10 mM Tris pH 8.0, 1mM EDTA with 100 μg RNase A (QIAGEN) was added to de-crosslinked DNA and incubated for 30 minutes at 37°C followed by addition of 5.0 μL of 20 mg/mL Proteinase K (QIAGEN) and incubation at 45°C for 30 minutes to remove RNA and proteins. DNA was purified with QIAGEN QIAquick purification columns and used for qPCR. De-crosslinked and purified input DNA (1 μL) was used for the Bioanalyzer assay to assess chromatin fragmentation.

### qPCR

Each pulldown was analyzed in quadruplicate using SYBR Green qPCR (Applied Biosystems, A46012). Input DNA was typically diluted 10x to reach similar concentration to ChIP DNA. Forward and reverse primers synthesized by Integrated DNA Technologies at 400 nM were combined with 2 μL DNA and 10 μL SYBR Green Mix in a 20 μL final reaction volume. Enzyme was activated at 95°C for 2 minutes before 40 cycles of denaturation and annealing (95°C for 5 s, 60°C for 30 s). The following primers were used (proximity of gRNA1 and 2 allowed the same primer pair): g1/2 Fwd-5’-TGGCTGTTTGTGGGTGGGTTT-3’, Rev-5’-GAAGCCCTCCCTATCACCGAA-3’; g3 Fwd-5’-GGAGTCAAGAAACTCCTGGGC-3’, Rev-5’-TCAAGGCAGCAGTCATCCCT-3’; *GAPDH* control Fwd-5’-TTGGCTACAGCAAGAGGGTG-3’, Rev-5’-GGGGAGATTCAGTGTGGTGG-3’. Enrichment was determined using the 2^-ΔΔCt^ method. Input DNA was normalized to 100% by subtracting the appropriate dilution factor from the raw Ct value (e.g. 9.966 for 10x dilution of 1% input [log_2_{1000}]). The Ct values for each individual gRNA from a single technical replicate were pooled (biological and technical replicates performed simultaneously in parallel for all gRNAs and NoG+ control). The degree of enrichment after ChIP was calculated by comparing the Ct of ChIP DNA for pooled gRNAs or NoG+ control normalized to the off-target *GAPDH* control with the Ct of adjusted input DNA normalized to off target *GAPDH*, i.e. ChIP(Ct[pooled gRNAs or NoG+] – Ct[*GAPDH*]) – adjusted input(Ct[pooled gRNAs or NoG+] – Ct[*GAPDH*]) = ΔΔCt. The fold change was calculated as 2^-ΔΔCt^. Fold change for biological and technical replicates were averaged and presented numerically as mean ± SEM and visually as a box and whisker plot.

### Proximity labeling

Cells were treated with 1 μg/mL doxycycline 21 hours before labeling and Caspex expression was confirmed by fluorescence of the GFP marker. Biotin tyramide phenol was diluted in external media from a 500 mM stock solution in DMSO to solubilize before being added to cells at a final concentration of 500 μM. After 30 minutes of incubation to allow time for the rate limiting step of biotin diffusion across membranes [169], hydrogen peroxide was diluted in media to 100 mM before addition to the cells at a final concentration of 1 mM to induce biotinylation. After 60 s of very gentle swirling, the media was decanted as quickly as possible and cells were washed three times with ice cold quenching solution (100 mM sodium azide, 100 mM sodium ascorbate in PBS), taking care to avoid dislodging the cells from the dish. Cells were washed with once ice cold PBS to help remove excess biotin phenol, transferred to 15 mL Falcon tubes, pelleted, flash frozen in liquid nitrogen, and stored at −80°C until use. For confirmation of proximity labeling, one 10-cm^2^ dish was used per biological replicate. For mass spectrometry studies, eight 15-cm^2^ dishes were used per replicate per condition.

### Western Blot

Labeled cell pellets were thawed at 4°C with gentle rotation in ice cold PBS. Pelleted cells were lysed in ice cold RIPA with freshly added protease and phosphatase inhibitors, DTT, and 10 mM sodium ascorbate and 10 mM sodium azide added to inhibit APEX2 to prevent excess labeling with any remaining adventitious biotin phenol. Lysate was sonicated on ice for three 10 s cycles. Forty micrograms of protein were denatured in NuPAGE LDS sample buffer (Invitrogen) supplemented with 50 mM DTT, followed by SDS-PAGE using 4-12% Bis-Tris WedgeWell Gel (Invitrogen) in MOPS running buffer (Invitrogen). Proteins were transferred to nitrocellulose membranes (0.2 μm, Bio-Rad) with a Trans-Blot Turbo Transfer System (Bio-Rad), blocked with 5% milk in TBST (30 min) and incubated with GAPDH Rabbit monoclonal RM114 antibody (Sigma SAB5600208) at 4°C overnight at a 1:2000 dilution. Following primary antibody incubation, membranes were washed three times with TBST, incubated with horseradish peroxidase-conjugated secondary anti-rabbit IgG (Cell Signaling 70745) and Streptavidin (Abcam AB279315) for 30 min (1:1000 and 1:10000 dilutions, respectively). Membranes were washed four times and signals were developed with chemiluminescence and imaged using the ChemiDoc Imaging system (Bio-Rad). Images were processed with the Fiji implementation of ImageJ [170]. Changes in contrast were applied to the entire blot.

### Enrichment of biotinylated proteins and on bead digestion

Cell pellets were thawed at 4°C with gentle rotation in cold PBS. Pelleted cells were lysed in 3 mL ice cold RIPA buffer with protease and phosphatase inhibitors, DTT, sodium ascorbate, and sodium azide added fresh before use. Lysate was sonicated as described above for ChIP. Typical sonication conditions resulted in an average size of ∼450 bp with 85-90% of chromatin between 200-1000 bp as assessed by the Bioanalyzer assay (**SF1c**). Protein concentration was determined by the Bradford assay at 595 nm (Abcam AB102535) – the addition of redox quenchers to prevent excess biotin labeling precludes use of the common BCA assay. For each condition, 500 μL of a Streptavidin M280 magnetic bead slurry (Dynabeads Invitrogen, 10 mg/mL) was utilized. The magnetic beads were washed twice with ice cold RIPA buffer. Lysates of equal protein amounts for each condition were incubated with pre-cleared beads for 120 minutes at room temperature. After capture of biotinylated proteins in the Falcon tube, beads were fully transferred to a 2mL Eppendorf protein lo-bind tube. Contaminants were removed by washing twice with 1 mL lysis buffer, once with 1M KCl, once with 100 mM Na_2_CO_3_, and twice with 6M GdCl in 50 mM triethylammonium bicarbonate (TEAB) (all ice cold). Proteins were denatured, reduced, and alkylated by incubation with 100 μL 6M GdCl, 10 mM TCEP, 40 mM chloroacetamide in 100 mM Tris pH 8.5 for five minutes at 95°C followed by fifty-five minutes at room temperature. Beads were washed four times with ice cold 50 mM TEAB to remove excess GdCl to prevent incomplete protease digestion. 1% of the bead mixture was removed to check capture efficiency via biotin elution (*vide infra*) before digestion. To a 20 μL slurry of bead-bound proteins in TEAB was added 400 ng (2 μL) of trypsin/LysC (Promega) [171] and the reaction was incubated at 37°C overnight. Digestion was quenched by the addition of 10% trifluoroacetic acid (TFA) to a final concentration of 1%. Supernatant was collected and the beads were washed with 25 μL of 1% TFA and supernatants were combined. Peptides were stored at −80°C until use. Remaining beads were eluted with biotin buffer to check digestion efficiency [42]. Briefly, beads were incubated with 50 μL 2x LDS loading buffer, 200 mM DTT, and 15 mM biotin for 15 minutes at 95°C two times. Combined supernatants were diluted 2x in LDS loading buffer and used for Western blot.

### TMT Labeling

Peptides were desalted with a Waters Oasis HLB Cartridge before labeling with TMTpro 18-plex reagent (Thermo) according to the manufacturer’s protocol. Briefly, TMT label reagents in anhydrous acetonitrile were added to each sample. The TMT isotopic labels were randomized across the different conditions according to the following scheme: g2 Rep1:126; g2 Rep2:127N; g1 Rep3:127C; g1 Rep1:128N; NoG+ Rep3:128C; NoG+ Rep1:129N; NoG-Rep1:129C; empty:130N; NoG+ Rep2:130C; NoG-Rep2:131N; g3 Rep3:131C; g3 Rep1:132N; g3 Rep2:132C; sample pool: 133N; g1 Rep2:133C; NoG-Rep3:134N; g2 Rep3:134C; empty:135N. The labeling reaction was allowed to proceed for 1 hr at room temperature before quenching with 5% hydroxylamine for 15 min. Samples were dried with a SpeedVac (Thermo) before resuspension in 100 mM ammonium formate for LC-MS analysis.

### Online 2D-LC and Data Acquisition via RTS-SPS-MS3

Data were collected with a Waters Acquity UPLC M-Class 2D-LC system directly connected to an Ascend Tribrid Mass Spectrometer (Thermo). A 12 multi-fraction gradient over 220 min was utilized with injection of 15 μg of peptides. The first dimension of separation at high pH across a BEH column consisted of buffer A (20 mM ammonium formate at pH 10) and buffer B (acetonitrile) using 12 discontinuous steps of buffer B at 10.8%, 13.1%, 14.9%, 16.7%, 17.7%, 18.9%, 19.9%, 20.4%, 22.2%, 25.8%, 28.9%, and 45% at a flow rate of 2 μL/min before transfer to a low pH C18 analytical column (75 μm ID/ 10 μM tip ID x 28 cm C18-AQ 1.8 μm resin). For each step, a 5 min isogradient of %B was used (**SF2**). TICs in SF2 were produced with the RaMS package in R [172]. In the second dimension, a linear gradient from 5% to 30% buffer B (0.1% formic acid in acetonitrile) in buffer A (0.1% formic acid in water) at a flow rate of 300 nL/min was used and directly injected to the mass spectrometer. The 3s duty cycle scan sequence began with an Orbitrap MS1 spectrum with the following parameters: resolution 120,000, scan range 400-1500 *m/z*, automatic gain control (AGC) target of 400,000 (100%), maximum injection time 50 ms, and RF lens 50% with a minimum of 6 points desired across the peak. Precursors were filtered using monoisotopic peak determination, charge state 2-7, and dynamic exclusion of 60 s with a ± 10 ppm tolerance excluding isotopes and different charge states. MS2 spectra were collected in the linear ion trap at a rapid scan rate with precursor fit of 50% in a 0.7 *m/z* window with a scan range of 400-1500 *m/z* (*mode auto*). Ions were fragmented with CID at a collision energy of 33% (activation time 10 ms, Q=0.25) with a maximum injection time of 40 ms and AGC target of 25,000 (250%). MS2 ions were selected by quadrupole across a 300-1500 *m/z* range for RTS-SPS analysis against the UniProt 2022 human protein database. A maximum of 4 peptides per protein were selected for RTS-SPS analysis. Enzyme specificity was set to Trypsin/P with a maximum of 1 missed cleavage and 35 ms maximum search time. A maximum of 2 variable modifications were allowed. Static modifications of cystine carbamidomethylation (57.0215 Da) and lysine and n-terminal TMTpro modification (304.2071 Da) plus variable methionine oxidation (15.9949 Da) were included for selection to MS3. Scoring thresholds for each charge state were set to Xcorr = 2.5(2+), 2.6(3+), 3.2(4+), and 3.2(5+) with dCn = 0.1 for all states. MS3 was measured in the Orbitrap at a resolution of 60,000 with fragmentation achieved via HCD at 65% collision energy. The scan range consisted of 100-500 *m/z* with a maximum injection time of 150 ms, AGC target of 100,000 (200%), and 8 precursors being isolated.

### Protein Quantification and Bioinformatic Analysis

Protein groups consistent with identified peptides were inferred with Proteome Discoverer 2.5 (Thermo). Raw files were searched against the UniProt 2022 human proteome database. Mass tolerance of 10 ppm was used for precursor ions and 0.6 Da for fragment ions. The search included cysteine carbamidomethylation (57.0215 Da) as a static modification. Peptide N-terminal and lysine TMTpro 18plex modification (304.2071 Da), protein N-terminal acetylation (42.0106 Da), and methionine oxidation (15.9949 Da) were set as dynamic modifications. Up to two missed cleavages were allowed for trypsin digestion. The peptide false discovery rate (FDR) was set as <1% using Percolator [173] with a reversed-sequence decoy database [174]. For protein identification, at least one peptide with a minimum 6 amino acid length was required. Grouped abundance for proteins from each TMT channel in **Supplementary Table S3** was used for bioinformatic analysis in Excel.

Grouped abundance of samples was first median normalized within each TMT channel, after which biological replicates were pooled together across TMT channels. Individual gRNA cell lines were treated as pseudo-technical replicates and pooled together for analysis. Raw quantification values approximated a binomial distribution before and after median normalization and were log_2_ transformed to approximate a normal distribution. The difference between log_2_ quantified conditions of interest for individual proteins was converted to a Z-score to enforce normality for statistical analysis, and a corresponding two-tailed *p*-value was calculated. The Z-score was calculated as x/σ, where x = difference in log_2_(median normalized protein quantification) and σ = standard error of the mean. Quantification error of the SEM for the log_2_ transformation was propagated via the delta method (d/dx log_2_(σ) = σ*(1/y*ln(2))), where y = median normalized quantification. Error of summation was propagated in the standard way (square root of the sum of squared errors). Proteins that were not quantified were excluded from analysis. Proteins that were not detected across all four conditions (NoG-, NoG+, pooled gRNAs, sample pool of all replicates) were excluded from analysis to prevent undefined ratios after manual inspection to assess potential biological significance. A contaminant list was created by comparing NoG-to the pooled labeling conditions (all gRNAs + NoG+) and proteins enriched by 1.2FC and Bonferroni adj. *p*<.05 were excluded from downstream analysis. Differences between pooled on-target conditions vs. NoG+ from the resulting list were then analyzed. Proteins with a FC>1.2 and a Storey-*q* value<.05 were considered enriched. Calculated *p*-values of zero were replaced with the lowest calculated *p*-value to prevent an undefined value in the volcano plot.

### Gene Ontology Analysis

Gene Ontology analysis was performed with DAVID, GORilla, and Panthedb. For each tool, the list of 6,039 inferred proteins was uploaded as a background list rather than using the whole human proteome to test for GO enrichment. Proteins enriched by FC>1.2 with (373) or without (775) the *q*-value filter were tested for enrichment of GO Biological Process, GO Cellular Component, and GO Molecular Function in each tool. For PantherdbGO, GO Complete terms were searched and Fisher’s exact test with Benjamini-Hochberg correction was used to determine significance. For DAVID, the top three Functional Annotation Clusters were recorded alongside the Functional Annotation Chart with Benjamini-Hochberg correction used for all comparisons. Benjamini-Hochberg correction was used for GORilla analysis in each category.

### Predicted Transcription Factor Binding

For TFBSPred, the first Transcription Start Site of FOXP2 (hg 38:7:114086327(+) - 114087534) was chosen for analysis (accessed August 2024). Every box was checked for lineage, tissue, karyotype, and sex. Every cell line was chosen for analysis and the default TFFM search threshold of 0.90 was used. Every predicted transcription factor present within the whole sequence was compared against our mass spectrometry results. For PROMO, *homo sapiens* was selected for the species and the same sequence analyzed by TFBSPred was used for input (accessed August 2024). All predicted transcription factors were recorded and compared to mass spectrometry data in **Supplementary Table S6**.

### Comparison with ENCODE ChIP-Seq Database

All published transcription factor and histone experiments that passed ENCODE data quality standards in the ChIP-Seq data matrix performed in HEK293 or HEK293T cells available at encodeproject.org were analyzed. The ENSCR experiment identifiers and associated laboratories for the 222 analyzed experiments are recorded in **Supplementary Table S7**. To determine if a factor binds to the *FOXP2* genomic locus, the web-based genome browser tool for each experiment was utilized. Only signals that were called as significant by ENCODE in the combined replicate experiments that reached the IDR threshold were considered as true hits – a significant signal in one or both replicates that did not persist in the combined experiments was discarded. Each experiment was analyzed at the following genomic loci (hg38 chr7): TSS1: 114084641-114087534; e330: 114411165-114418591; upstream intergenic region: 114075920-114084640; *FOXP2* gene body: 114086327-114693773. Any peaks that passed the IDR threshold within the analyzed region were recorded as a binary Yes/No in **Supplementary Table S7**. Multiple significant peaks within a defined analyzed region were individually counted for the n>3 condition.

### Statistical Analysis

To assess statistical significance of enrichment values for ChIP-qPCR and expression changes for siRNA, a two-sided *t*-test was performed. To determine enrichment of transcription factors and components of the spliceosome, the χ^2^ test was performed on pairwise comparisons after creating a contingency table. For protein quantification, the *p*-value corresponding to the Z-score calculated for difference in protein quantification between states of interest was calculated. To correct for multiple hypothesis testing across the thousands of quantified proteins, Bonferroni or Storey methods were used as indicated in the text. For Bonferroni correction, the significance threshold was set at α/n, where α = uncorrected significance threshold (set to .05) and n = number of comparisons. Storey correction is a modification of Benjamini-Hochberg correction where the proportion of null hypotheses is estimated to be less conservative in calling significance. The results from the n statistical tests were listed in ascending order of *p*-value and assigned a rank order of 1,2,…,n. The proportion of null hypotheses, π_0_, was estimated according to the formula π_0_ = (#*p*-values>λ)/(n*(1-λ)), where λ = 0.4 as estimated from the flat portion of the *p*-value histogram. The value λ represents uniform distribution of truly null values considering the fact that truly significant values will skew towards lower *p*-values. The *q*-value for element *i* was calculated as the lower of (π_0_*n\**p*-value*_i_*)/rank*_i_* and *q*-value(*i*+1).

## Data Availability

Mass spectrometry data described in this communication have been deposited to the MassIVE repository partner of the ProteomeXchange Consortium available at massive.ucsd.edu with the identifiers <u>MSV000098217</u> (MassIVE) and <u>PXD065063</u> (ProteomeXchange) with the password Language. Experiment identifiers and contributing labs for the 222 ENCODE ChIP-Seq experiments utilized in this study are enumerated in **Table S7**. Data for IRF2BP2 knockdown and ChIP-Seq were obtained through the NCBI GEO with accession identifier <u>GSE124636</u>.

## Description of Supplemental Attachments

**Supplemental figures: SF1** Preparing Chromatin for Analysis; **SF2** Total Ion Chromatograms from Each High pH Fraction from 2D-LC.

**Table S1** Peptide Spectral Matches from Proteome Discoverer Analysis

**Table S2** Peptide Groups from Proteome Discoverer Analysis

**Table S3** Proteins Inferred from Proteome Discoverer Analysis

**Table S4** Gene Ontology Analysis for Enriched Proteins with and without *q*-value Filter

**Table S5** Comparison of Human Transcription Factors and Spliceosome Components to Proteins Enriched with and without *q*-value Filter

**Table S6** Predicted Transcription Factors Recognizing *FOXP2* TSS1 from TFBSPred and PROMO

**Table S7** ENCODE ChIP-Seq Results at *FOXP2* and Comparison to Mass Spectrometry Results

